# Evidence of Lactate Shuttling in the Human Brain using Hyperpolarized ^13^C-MRI

**DOI:** 10.1101/2023.01.13.523957

**Authors:** Biranavan Uthayakumar, Hany Soliman, Albert P. Chen, Nadia Bragagnolo, Ruby Endre, William J. Perks, Nathan Ma, Chris Heyn, Charles H. Cunningham

## Abstract

**Purpose:** To test the hypothesis that lactate shuttling contributes to the ^13^C-lactate and ^13^C-bicarbonate signal observed in the awake human brain using hyperpolarized ^13^C MRI.

**Methods:** Healthy human volunteers (n = 6) were scanned twice using hyperpolarized ^13^C-MRI, with reduced radiofrequency saturation of ^13^C-lactate on one set of scans. ^13^C-lactate, ^13^C-bicarbonate, and ^13^C-pyruvate signals for 132 brain regions across each set of scans were compared using a clustered Wilcoxon sum rank test.

**Results:** Reduced ^13^C-lactate radiofrequency saturation resulted in a significantly greater ^13^C-bicarbonate signal (*p* = 0.04). These changes were observed across the majority of brain regions.

**Conclusion:** Radiofrequency saturation of ^13^C-lactate leads to a decrease in ^13^C-bicarbonate signal, demonstrating that the ^13^C-lactate generated from the injected ^13^C-pyruvate is being converted back to ^13^C-pyruvate and oxidized throughout the human brain.

## Introduction

It is well known that glucose is the primary source of energy in the brain, but mounting evidence suggests that at least some of this glucose is first converted to lactate and shuttled between cellular compartments before ultimately being converted back to pyruvate and oxidized in the TCA cycle (1, 2). In this study, the hypothesis that “lactate shuttling” contributes to the ^13^C-lactate and ^13^C-bicarbonate signal observed in the awake human brain is tested using hyperpolarized ^13^C MRI (HP^13^C-MRI).

HP^13^C-MRI is a methodology for metabolic profiling *in vivo*. Following the intravenous injection of ^13^C-pyruvate, the downstream metabolites ^13^C-lactate and ^13^C-bicarbonate can be readily imaged in the human brain (3, 4, 5, 6). Signal from ^13^C-lactate shows intracelluar conversion of ^13^C-pyruvate via the enzyme lactate dehydrogenase (LDH) and is higher with increased local rate of lactate production and/or increased local lactate pool size.

The ^13^C-bicarbonate signal represents bicarbonate created when ^13^C-pyruvate is converted to acetyl-CoA on the mitochondrial membrane, with the acetyl-CoA going on to be oxidized in the TCA cycle. Thus ^13^C-bicarbonate signal reflects the flux of ^13^C-pyruvate into mitochondrial oxidation.

However, if lactate shuttling is occurring at a significant level in the human brain, some fraction of the ^13^C-bicarbonate could be formed from ^13^C-lactate that has been created in one cellular compartment, shuttled to a separate physical environment, and then converted back to ^13^C-pyruvate and consumed in the TCA cycle in a separate compartment (see Fig. 1). The astrocyte-neuron lactate shuttle (ANLS) theory (1) proposes that lactate is predominantly produced in one cellular compartment (astrocytes) and then shuttled to a separate compartment (neurons) where it is converted back to pyruvate and used in mitochondrial oxidative phosphorylation. Evidence of ^13^C-lactate shuttling in the rat brain was previously demonstrated (7), with ^13^C-bicarbonate signal in the brain reduced when RF saturation was applied to the ^13^C-lactate pool. In this study, the hypothesis that^13^C-lactate shuttling is occurring in the human brain was tested using two consecutive HP ^13^C-MRI scans with two different flip angles (46° vs 80°) applied when imaging ^13^C-lactate.

**Figure 1:**
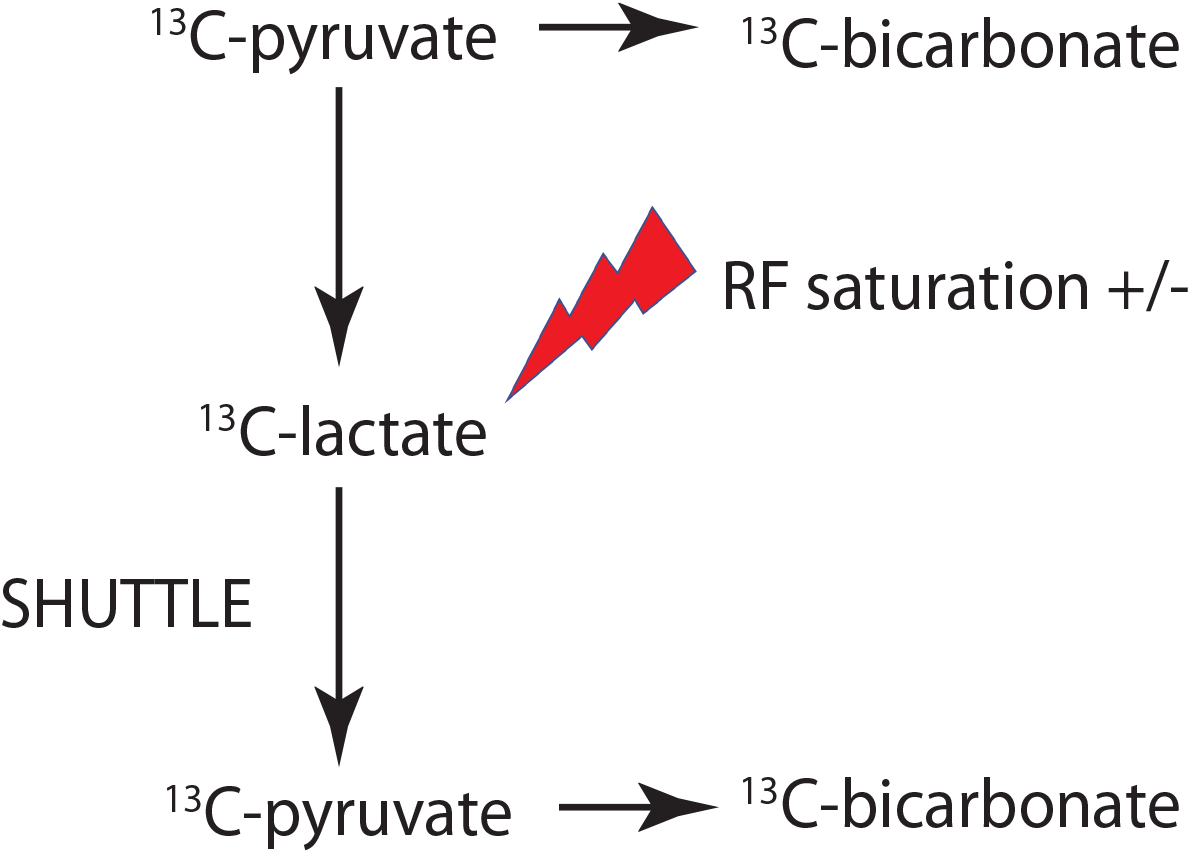
Experiment schematic, showing ^13^C-pyruvate taking two possible pathways to result in ^13^C-bicarbonate. Varying degrees of RF saturation are applied to the ^13^C-lactate lactate pool to test whether increased saturation of ^13^C-lactate causes a decrease in ^13^C-bicarbonate signal, indicating that ^13^C-lactate is being used in oxidative metabolism.

## Materials and Methods

Study participants (N=6) were between the ages of 23 and 48 and screened for cognitive impairment using the Montreal Cognitive Assessment (MoCA) (8). The experiment was conducted under a protocol approved by the Research Ethics Board of Sunnybrook Health Sciences Centre and regulated by Health Canada under a Clinical Trial Application. All participants gave written informed consent.

### Data Acquisition

Each dose was compounded from a 1.47 g sample of [1-^13^C]pyruvic acid (Sigma Aldrich, St. Louis, MO) using a SPINLab polarizer system (GE Healthcare, Waukesha, WI). A 20 gauge intravenous catheter was placed in the forearm of each participant prior to scanning for injection of the hyperpolarized [1-^13^C]pyruvate solution.

Imaging was performed using a GE MR750 3.0T MRI scanner (GE Healthcare, Waukesha, WI), and a ^13^C birdcage head coil built in-house. A spectral-spatial excitation and 3D dual-echo echo-planar imaging readout were used to acquire images of ^13^C-lactate, ^13^C-bicarbonate, and ^13^C-pyruvate with a temporal resolution of 5 seconds and spatial resolution of 1.5 cm (9). Each volumetric ^13^C-metabolite image was acquired at 5 second intervals over the course of a sixty second acquisition window.

The ^13^C scan was performed twice for each participant, with a half-hour wait time between scans. For one of the scans, a reduced flip angle was used for excitation of the ^13^C-lactate resonance, in order to reduce the saturation of the ^13^C-lactate pool, while the flip angle for ^13^C-bicarbonate and ^13^C-pyruvate remained the same for both scans.

The net flip angle was 80° for ^13^C-lactate, resulting from a 20° spectrally-selective excitation pulse applied 24 times for the 24 phase encodes in the slice direction, 80° for ^13^C-bicarbonate, and 11° for ^13^ C-pyruvate in the condition designated the ‘lactate-80°’ scan. The ‘lactate-46°’ scan was acquired with a net flip angle of 46° excitation of lactate (24 applications of a 10° excitation pulse), with the flip angles for the other ^13^C-metabolites held constant. The first three participants (volunteers 1-3) had the lactate-46° scan done as the second scan, while the latter three scans had the lactate-46° done as the first scan.

During the half-hour wait time between the two ^13^C-metabolite scans, the ^13^C head coil was interchanged with an 8-channel ^1^H neurovascular array (Invivo Inc, Pewaukee, WI) for anatomical imaging. Anatomical images were acquired using axial fast spoiled gradient echo images (FOV 25.6×25.6 cm^2^, 1 mm isotropic resolution, TR 7.6 ms, TE 2.9 ms, flip angle 11°). Al 1 ^13^C-images were reconstructed and resampled to the resolution of the anatomical images in Matlab (10) before being saved in DICOM format.

### Data Analysis

For each subject, the metabolite images from the full 60 second acquisition window were summed to produce time-integrated ^13^C images. The T1-weighted anatomical images were parcellated into the 132 brain regions in the BrainCOLOR labelling protocol (11) using the Spatially Localized Atlas Network Tiles (SLANT) method (12). The parcellation maps were then used to compute mean ^13^C-pyruvate, ^13^C-lactate and ^13^C-bicarbonate signal for each region and subject using the mrisegstats software (http://nmr.mgh.harvard.edu). These are referred to below as regional metabolite signal values and are in measured arbitrary units but are comparable between subjects and conditions despite additional sources of variance such as slightly differing ^13^C polarization levels.

In order to test for a significant difference in the regional ^13^C-metabolite signal between the two conditions, a clustered Wilcoxon signed rank test was used (13). The paired ^13^C-bicarbonate signals (each pair for the lactate-46° and lactate-80° conditions, for all of the atlas regions from a each individual participant were designated as a cluster and the clusrank package (14) R version 3-3 (15)) was used to perform the Wilcoxon signed-rank test combining the data from the 6 participants. This was also performed for ^13^C-lactate and ^13^C-pyruvate signal.

To assess the relationship between the regional ^13^C-metabolite signal level and the regional percentage change between the lactate-46° and lactate-80° conditions, Spearman’s rank correlations were computed, using the mean of each brain region across subjects. The false-discovery rate method (16) was used to set the threshold for significance at p = 0.013, accounting for the 15 correlations performed. The correlations are summarized in Fig. 6.

To rule out the possibility of differences in ^13^C-pyruvate polarization being the primary cause of differences between the two conditions, a t-test was performed between the polarization levels for the two conditions, as well as for the time from dissolution to injection (N=6).

## Results

The increase in ^13^C-bicarbonate signal across regions is apparent in the example images from one participants in Fig. 2 and in the boxplots showing all of the regions for the two conditions in Fig. 3.

**Figure 2:**
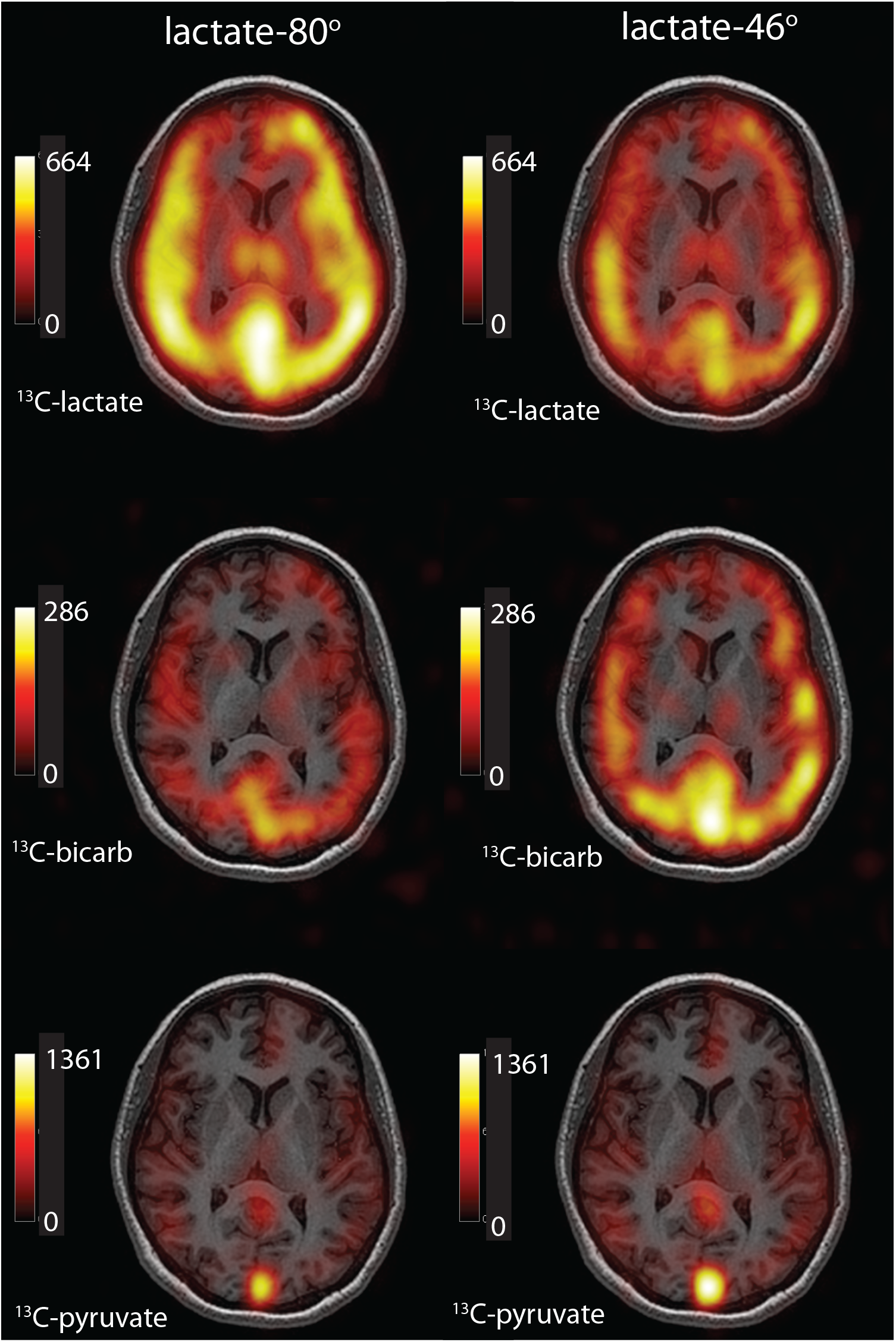
Example images from the lactate-80° (left) and lactate-46° (right) conditions. All images are from the same participant. The ^13^C-lactate images (upper row) show that ^13^C-lactate signal (colour overlay) is reduced in the lactate-46° condition, as expected. Conversely, the ^13^C-bicarbonate signal (middle row) is increased in the lactate-46° condition. The ^13^C-pyruvate signal (bottom row) shows increased signal in the sagittal sinus in the lactate-46° condition.

**Figure 3:**
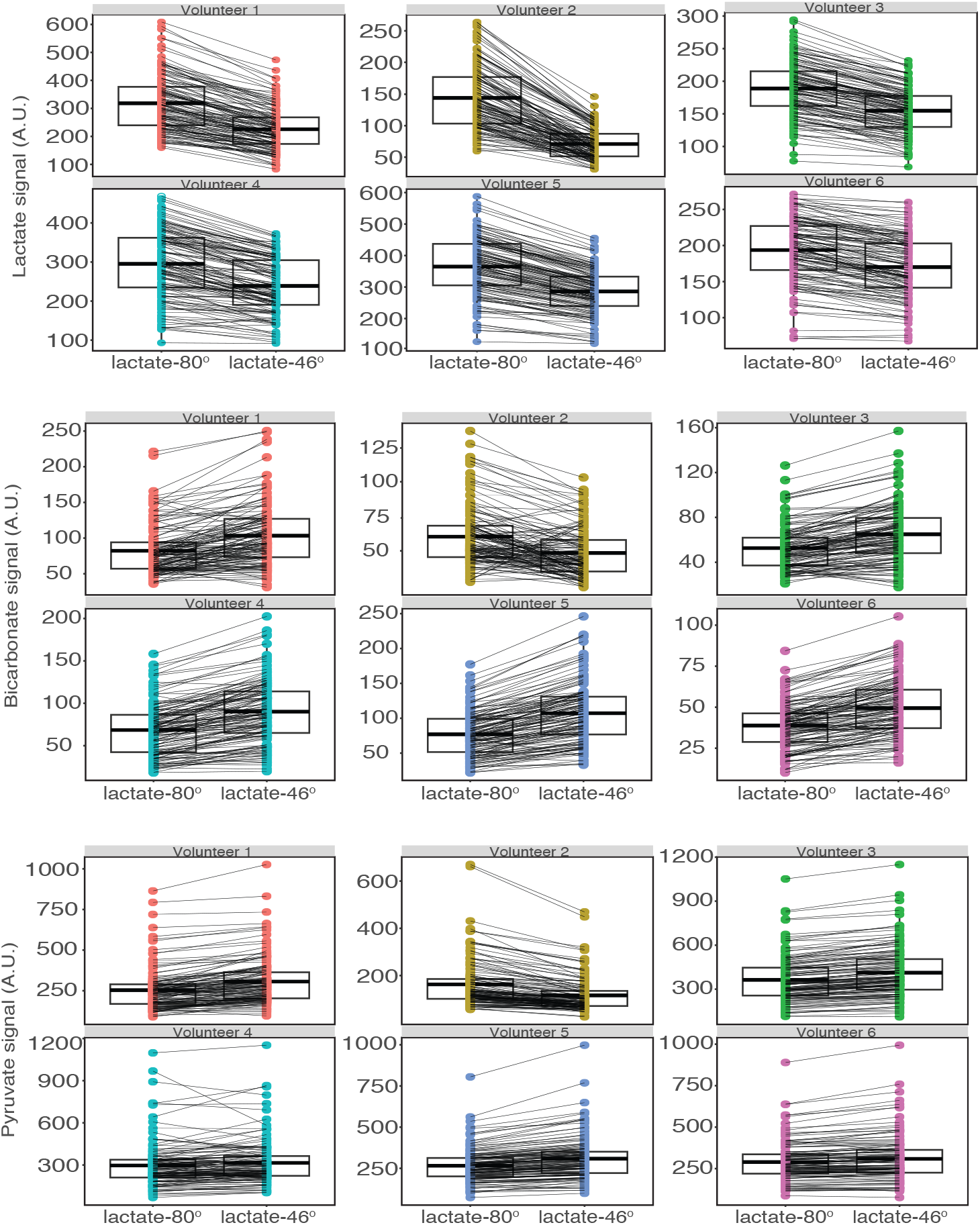
Boxplots of mean regional ^13^C-lactate, ^13^C-bicarbonate, and ^13^C-pyruvate signal for each of the six participants. The boxes in each panel shows the regional ^13^C-metabolite signal from the lactate-80° (left) and lactate-46° (right) conditions. Lines connect the same brain region between the two different conditions for each participant.

The clustered Wilcoxon signed rank tests showed a significant difference between conditions for ^13^C-bicarbonate (*p* = 0.04, Z = 1.783), supporting the main hypothesis that decreased RF saturation of the ^13^C-lactate pool increases the ^13^C-bicarbronate signal. The ^13^C-pyruvate signal also showed an increase although this did not reach significance (*p* = 0.08, Z = 1.4). For ^13^C-lactate, the different flip angles used in the two conditions gave the largest and most significant change (decrease) in signal (*p* = 0.01, Z = −2.3), as was expected due to the reduction in flip angle.

The mean percentage change between the two conditions was −25% ±7% for ^13^C-lactate, +21% ±5% for ^13^C-bicarbonate and 6% ± 6% for ^13^C-pyruvate. A positive percentage change indicates higher ^13^C-metabolite signal for the lactate-46° condition relative to the lactate-80° condition. The ratio of ^13^C-bicarbonate signal from the lactate-80° scan to ^13^C-bicarbonate signal from the lactate-46° scan is plotted for each of the 132 brain regions and plotted in Fig. 4. Each box is composed of the N = 6 ratios for each brain region.

**Figure 4:**
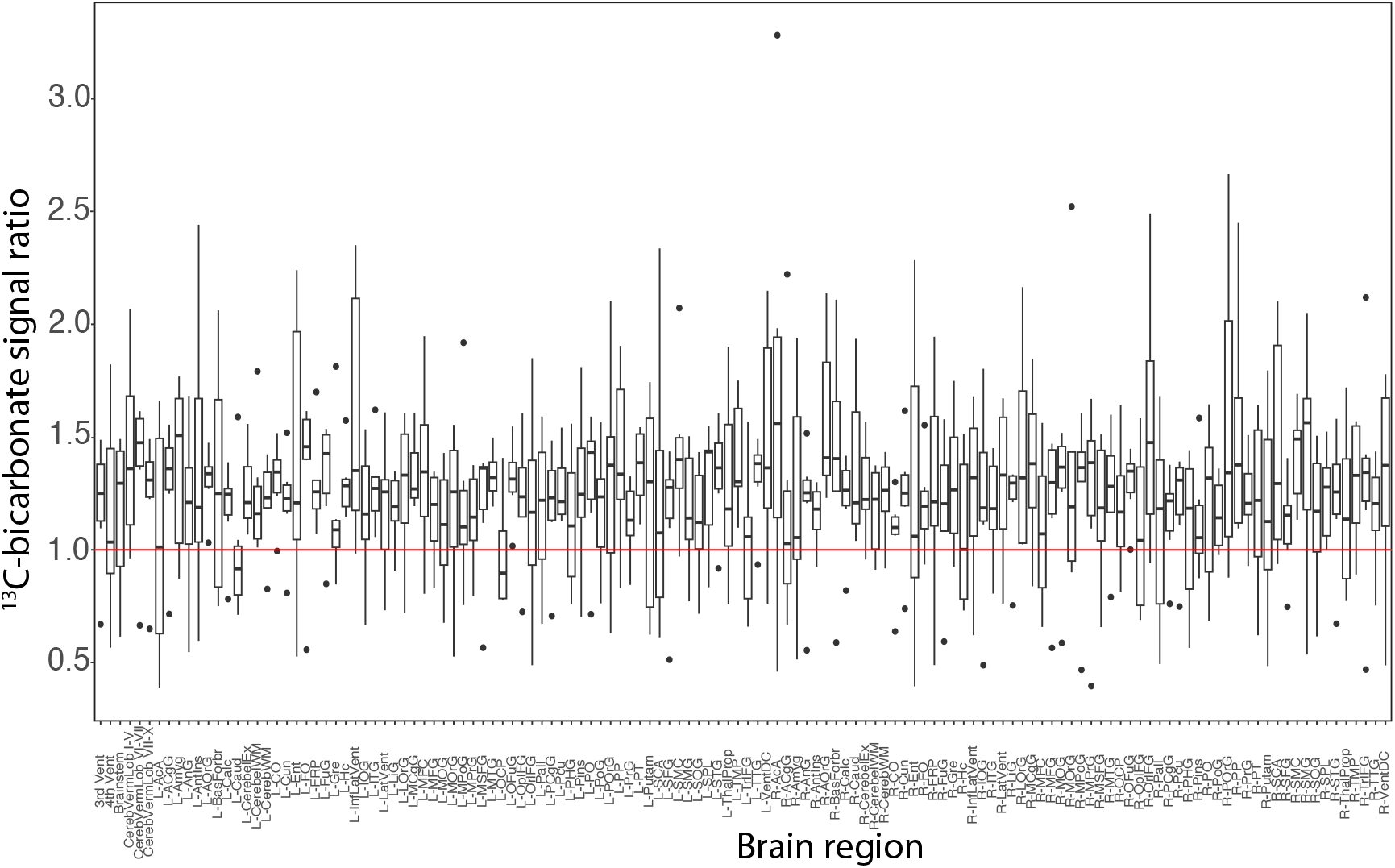
^13^C-bicarbonate signal measured in the lactate-80° condition divided by ^13^C-bicarbonate signal measured in the lactate-46° condition for each of the 132 brain regions. Each boxplot comprises the N=6 ratios for that region. The red line shows unity, with equal signal in the two conditions.

The majority of brain regions have mean values above 1.0 (red line), which indicates increased ^13^C-bicarbonate signal with reduced ^13^C-lactate RF saturation. A similar effect was observed for ^13^C-pyruvate signal (Fig. 5, with the ratio for most regions above 1.0, indicating increased ^13^C-pyruvate signal with reduced RF saturation of ^13^C-lactate. Both of these results are consistent with the SHUTTLE pathway in Fig. 1 being a significant source of ^13^C-bicarbonate signal in the human brain.

**Figure 5:**
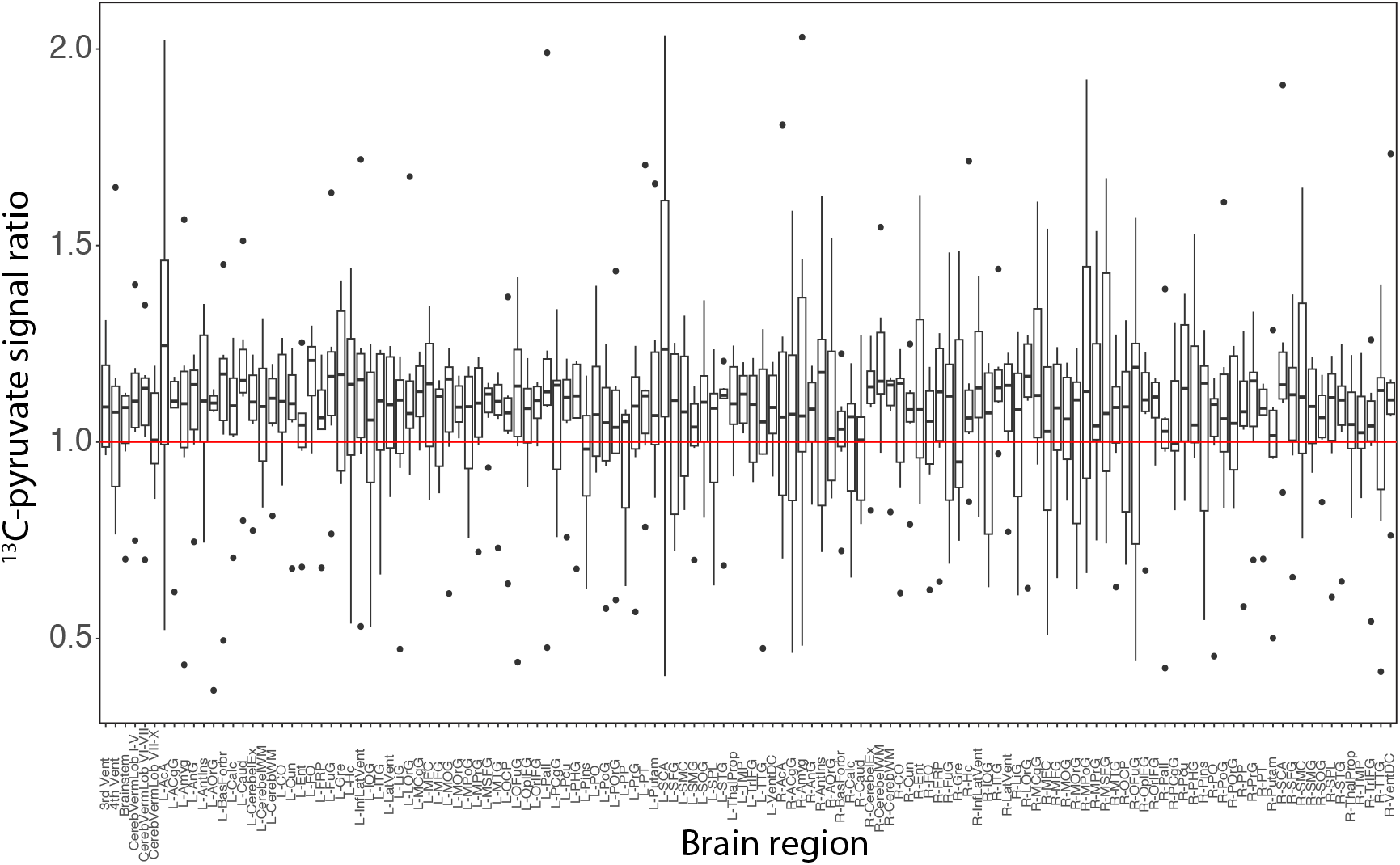
^13^C-pyruvate signal measured in the lactate-80° condition divided by ^13^C-pyruvate signal measured in the lactate-46° condition for each of the 132 brain regions. Each boxplot comprises the N=6 ratios for that region. The red line shows unity, with equal signal in the two conditions.

**Figure 6:**
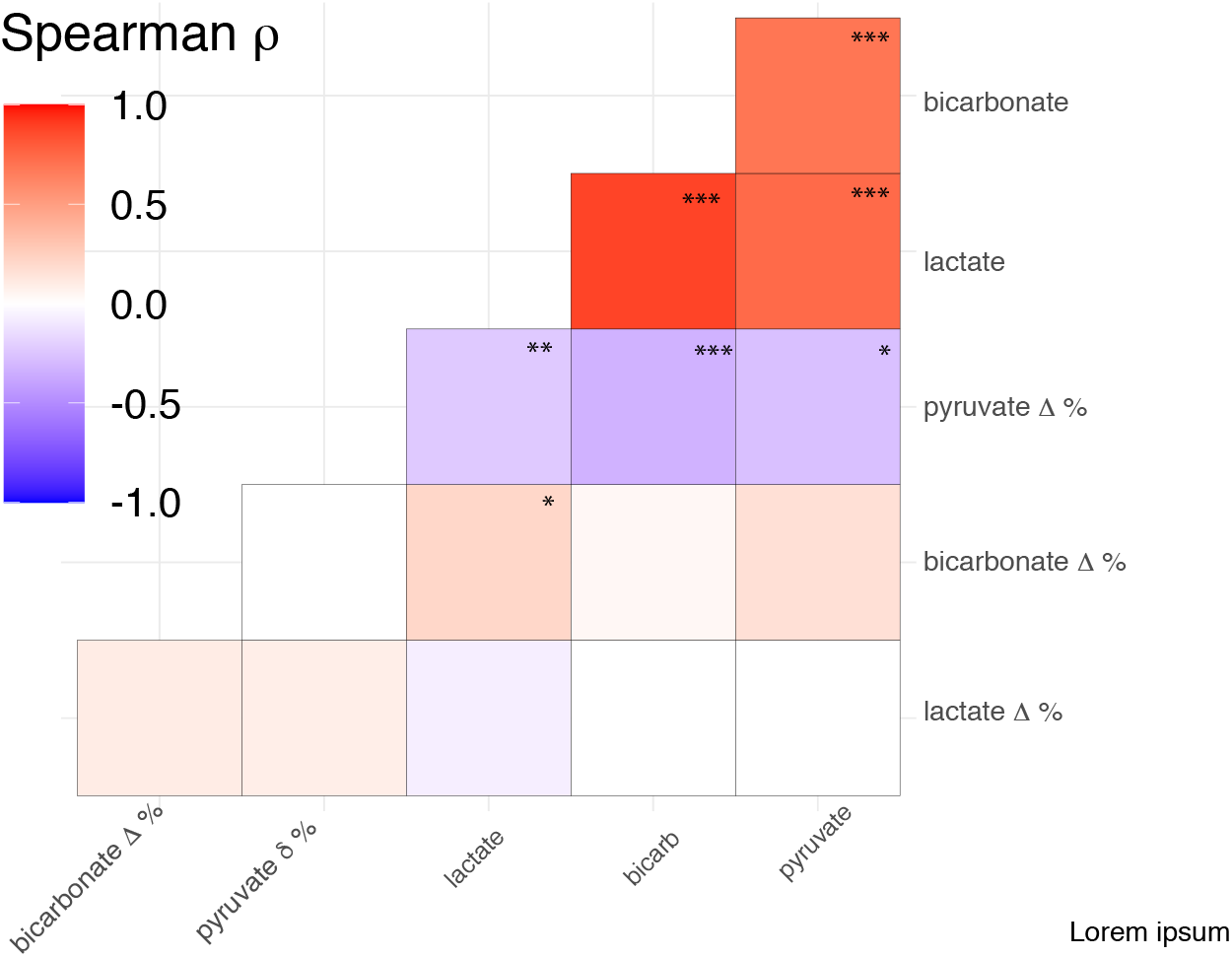
Correlation matrix of the percent changes in each ^13^C-metabolite signal between conditions as a function of each ^13^C-metabolite (\**p* ≤ 0.05, \*\**p* ≤ 0.01 \*\*\**p* < 0.001)

The percentage change (denoted Δ %) in regional ^13^C-bicarbonate signal between the lactate-80° and lactate-46° conditions are plotted vs. the mean regional signal (from the lactate-80° condition) for each of the three ^13^C-metabolites in Fig. 7. There was a weak but significant positive correlation between the percentage change in ^13^C-bicarbonate signal and the mean regional ^13^C-lactate signal, meaning that brain regions that produced more ^13^C-lactate were associated with increased shuttling and conversion of ^13^C-lactate back to ^13^C-pyruvate and entering mitochondria, resulting in ^13^C-bicarbonate.

**Figure 7:**
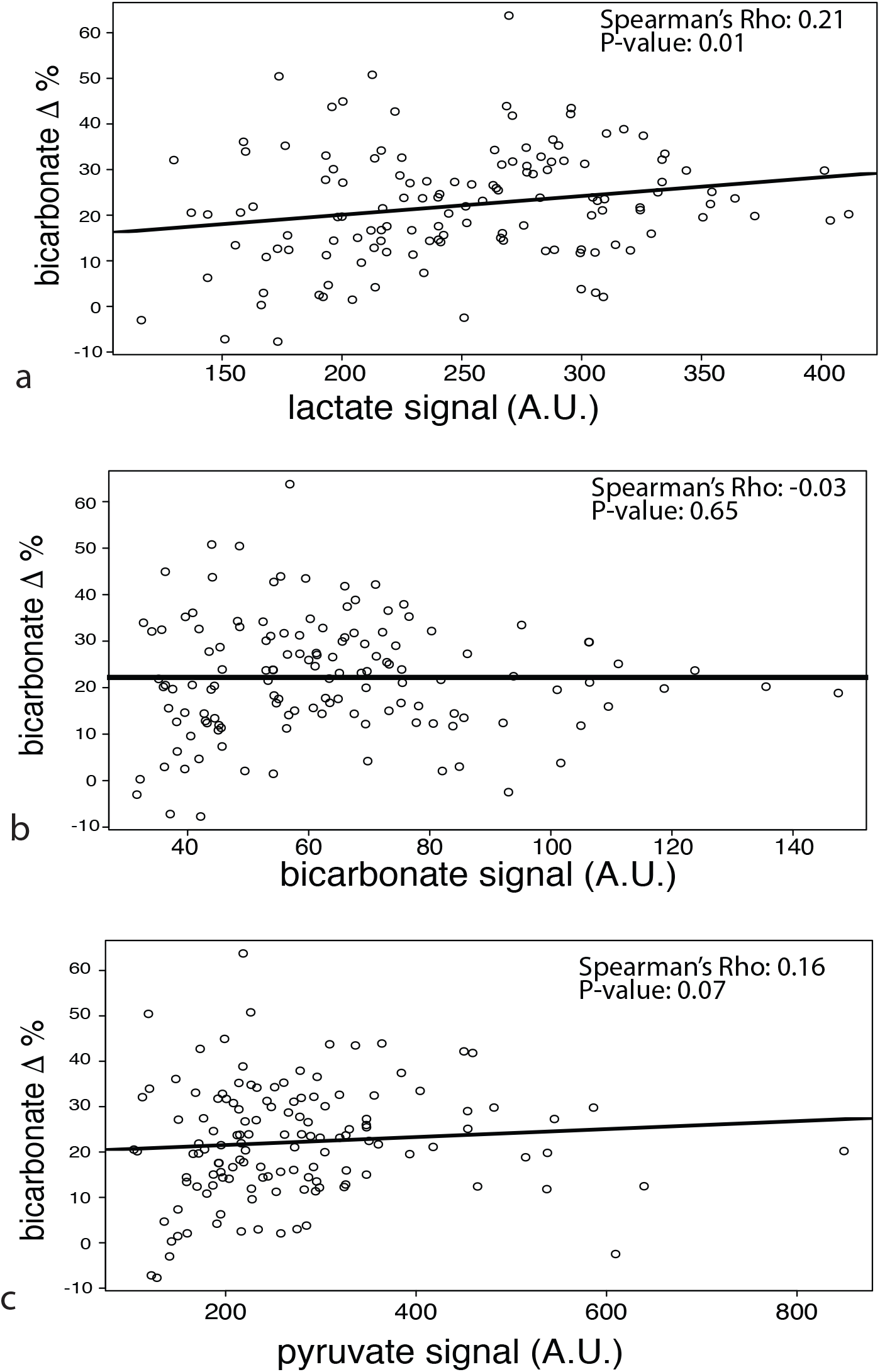
The percentage change (Δ %) in regional ^13^C-bicarbonate signal between the two conditions plotted vs mean regional ^13^C-metabolite signal. A significant positive correlation is seen in (a). Each point is a particular brain region averaged across the six healthy volunteers. The Spearman correlation coefficient and corresponding p-value are listed in each plot.

Conversely, the percentage change in ^13^C-pyruvate signal between the lactate-80° and lactate-46° conditions showed a weak but significant *negative* correlation with the regional ^13^C-metabolites signal (see Fig. 8). The opposing correlations observed for Δ% ^13^C-pyruvate and Δ % ^13^C-bicarbonate may be explained by the hypothesis that regions with lower ^13^C-pyruvate uptake contain a slightly higher fraction of ^13^C-lactate-derived ^13^C-pyruvate simply because the overall ^13^C-enriched pool size is smaller.

**Figure 8:**
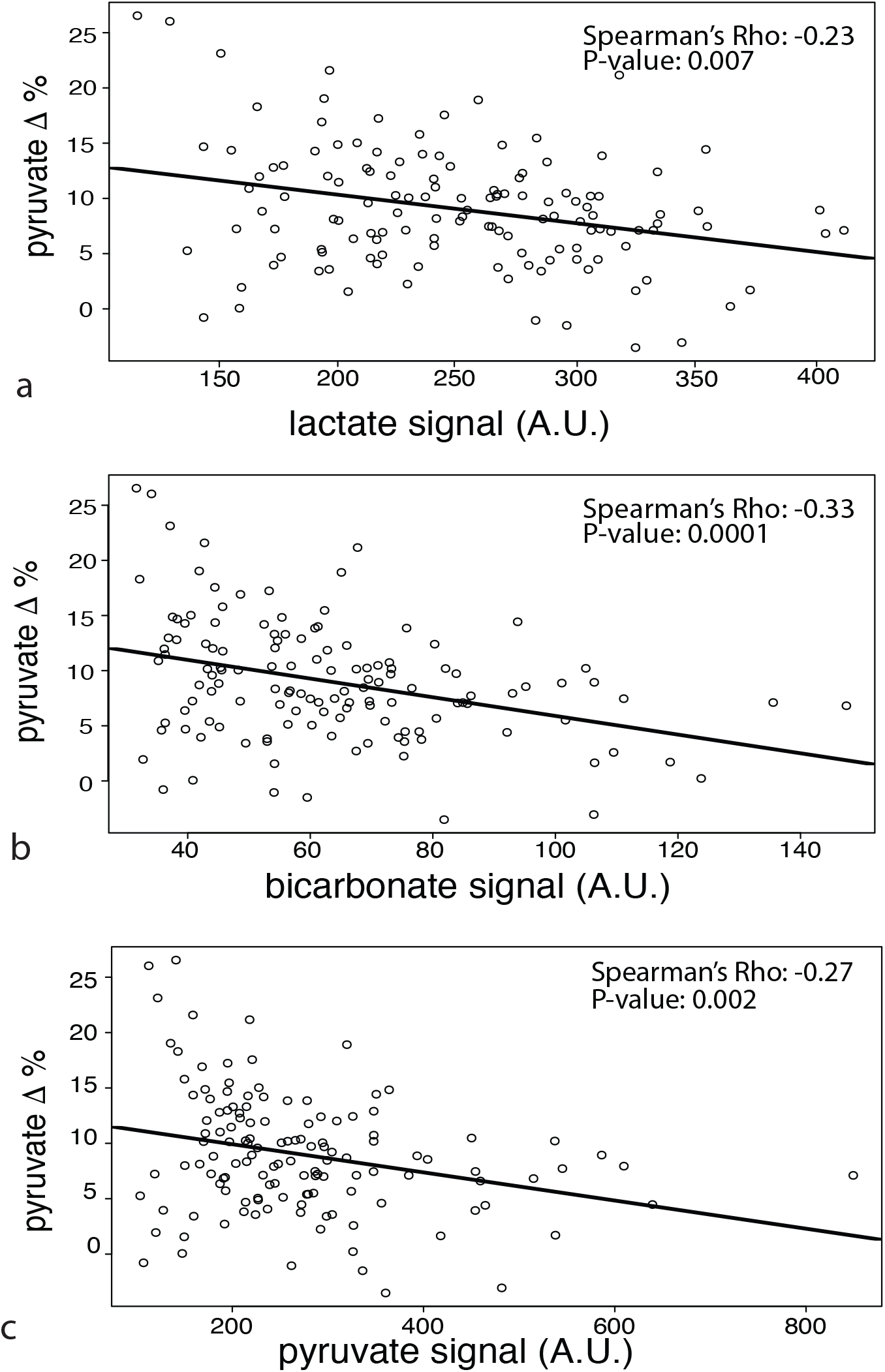
The percentage change (Δ %) in regional ^13^C-pyruvate signal between the two conditions plotted vs mean regional ^13^C-metabolite signal. All three plots show a significant negative correlation. Each point is a particular brain region averaged across the six healthy volunteers. The Spearman correlation coefficient and corresponding p-value are listed in each plot.

Supplementary Tab. 1 shows the liquid-state ^13^C polarization readouts from the qualitycontrol module on the SPINLab polarizer immediately after dissolution, as well as time-to-injection after dissolution. The t-tests comparing the polarization and time-to-injection for the lactate-80° vs. the lactate-4O° scans were both insignificant (*p* = 0.95 and *p =* 0.77, respectively).

## Discussion

Previous studies have shown evidence of lactate shuttling in the human brain using microdialysis and magnetic resonance spectroscopy (17, 18). In this study, it was shown that reduced RF saturation of the ^13^C-lactate pool resulted in increased ^13^C-bicarbonate signal. This result is in agreement with two prior studies in animals (7, 19) as well as a human participant (19).

The results suggest that ^13^C-lactate is converted into acetyl-CoA throughout the whole brain resulting in ^13^C-bicarbonate. An increase in ^13^C-pyruvate signal in the lactate-46° condition compared to the lactate-80° condition was also observed. This agrees with the model shown in Fig. 1 wherein ^13^C-lactate is converted back to ^13^C-pyruvate prior to conversion to acetyl-CoA via pyruvate dehydrogenase (PDH). The increase in both ^13^C-pyruvate and ^13^C-bicarbonate under the lactate-46° condition was observed across almost all 132 brain regions assessed. Differences between brain regions were not assessed due to the limited sample size. However, differential expression patterns of the enzymes LDH and PDH between brain regions, as well as the expression of regulating factors like pyruvate dehydrogenase kinase (PDK), which inactivates PDH, can be seen in maps of single-cell RNA-sequencing from the brain (20).

There were study limitations that should be addressed. The course 1.5 cm isotropic spatial resolution caused partial volume effects that likely contributed to the variance observed, although such an effects would not cause a systematic bias between the two conditions. Secondly, raw ^13^C signal values were used in the analysis without normalizing for any difference in polarization or substrate transit time between subsequent injections. However, any effect of differing polarization between the two conditions was mitigated by varying the temporal ordering of the two scans. Lastly, this experiment showed only that ^13^C-lactate is being converted back to ^13^C-pyruvate and being used in mitochondria, which is consistent with lactate shuttling, but it did not provide any direct evidence about the compartments involved. Adding diffusion weighting to future experiments may shed more light on this.

## Conclusion

This study provided evidence from the human brain that lactate shuttling is occurring in the human brain at a level that is readily observed with hyperpolarized ^13^C MRI. Injected ^13^C-pyruvate is converted to ^13^C-lactate for some time, and then converted back to ^13^C-pyruvate and then to acetyl-CoA resulting in ^13^C-bicarbonate. It remains to be shown whether the observed oxidative metabolism of lactate is occurring in neurons or glial cells and future experiments with diffusion weighting and other pulse-sequence manipulations may shed light on this question.

## Figures and Tables

**Table 1:**
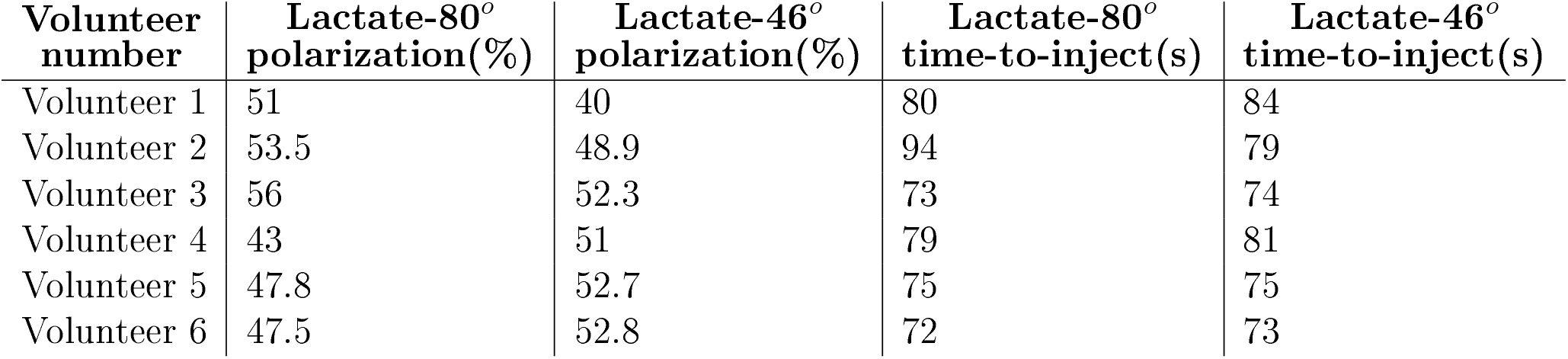
The polarization recorded from the quality-control module on the SPINLab polarizer and the time-to-injection for each of scan.

| **Volunteer**<br>**number** | **Lactate-80°**<br>**polarization(%)** | **Lactate-46°**<br>**polarization(%)** | **Lactate-80°**<br>**time-to-inject(s)** | **Lactate-46°**<br>**time-to-inject(s)** |
| --- | --- | --- | --- | --- |
| Volunteer 1 | 51 | 40 | 80 | 84 |
| Volunteer 2 | 53.5 | 48.9 | 94 | 79 |
| Volunteer 3 | 56 | 52.3 | 73 | 74 |
| Volunteer 4 | 43 | 51 | 79 | 81 |
| Volunteer 5 | 47.8 | 52.7 | 75 | 75 |
| Volunteer 6 | 47.5 | 52.8 | 72 | 73 |

## Abbreviations

ANLS: Astrocyte-neuron lactate shuttle
HP^13^C: hyperpolarized ^13^C
RF: radiofrequency
TCA: Tricarboxylic acid

